# Extensive MEG time-series phenotyping unveils neural markers predictive of age

**DOI:** 10.1101/2024.05.09.593348

**Authors:** Christina Stier, Elio Balestrieri, Jana Fehring, Niels K. Focke, Andreas Wollbrink, Udo Dannlowski, Joachim Gross

## Abstract

Understanding the evolving dynamics of the brain throughout life is pivotal for anticipating and evaluating individual health. While previous research has described age effects on spectral properties of neural signals, it remains unclear which ones are most indicative of age-related processes. This study addresses this gap by analyzing resting-state data obtained from magnetoencephalography in 350 adults (18-88 years). We employed advanced time-series analysis at the brain region level and machine learning to predict age. While traditional spectral features achieved low to moderate accuracy, over a hundred novel time-series features proved superior. Notably, temporal autocorrelation emerged as the most robust predictor of age. Distinct patterns of autocorrelation within the visual and temporal cortex were most informative, offering a versatile measure of age-related signal changes for comprehensive health assessments based on brain activity.

## 1. Introduction

Development and aging are complex processes encompassing profound structural and functional brain changes. Efforts to delineate linear and non-linear age effects on neurophysiological activity using large-scale datasets have increased in recent years. Notable changes of spectral characteristics during rest have been described in distinct brain regions, spanning childhood and adolescence (for example ^1,2^), and a broad age range during adulthood ^3–5^. These studies have been seminal in developing a deeper understanding of M/EEG measures widely applied in clinical and cognitive neurosciences. A common and early observation is that low-frequency power decreases and high-frequency power increases linearly with age ^6–10^, with the most pronounced changes in frontal and temporal areas of the brain ^2–4,11^. Studies on the brain’s functional connectome are less frequent and consistent partly due to the various connectivity measures used, relying on either amplitude or phase information of the signals. During adolescence, connectivity in the theta band seems to develop along an anterior-posterior axis such that decoupling between association areas is linked to cognitive control ^1^. From childhood through middle age, age-related changes have also been observed in the gamma range^2^. From early adulthood to old age, connectivity primarily changes in visual areas ^9^, partially irrespective of the frequency range ^4^. Depending on the age range investigated, power or connectivity measures showed non-linear age trajectories ^2^. During adulthood, for example, this was mostly the case in the delta and beta-gamma range and in regions that tend to be transmodal ^3,4,7^. While these studies aim to associate age with brain phenotypes and sometimes with behavioral phenotypes, they tend to be descriptive, thus limiting the generalizability of findings to other populations and clinical translation ^12^. Association studies are prone to overfitting such that a model usually fits the investigated sample better than a new, independent dataset. Predictive modeling aims to overcome these limitations, for example through cross-validation that uses subsets of the sample for iterative training and testing of the model ^13^.

In the last decade, openly accessible neuroimaging datasets have enabled predictive modeling using large sample sizes, mostly focusing on MRI-derived features in the context of development and aging (for an overview see ^14^). Abnormal brain aging, quantified by the difference between chronological age and predicted age of an individual, has been observed in several neurological and psychiatric disorders ^15–19^, linked to cognition ^20^ and physical health ^21,22^. These so-called brain age gaps likely underlie genetic influence ^15,23^ and represent specific biophysical processes ^24^, making it a suitable paradigm to study the interaction between normal aging and pathology ^25^. Notably, previous findings suggest that neural activity in the millisecond range improves brain age predictions derived from MRI ^26,27^. Engemann and colleagues report that electrophysiological and BOLD signals helped predict distinct cases of age using machine learning, yielding additive effects of the modalities ^27^. Moreover, age effects on fMRI and M/EEG signals have been complementary rather than redundant ^28^, urging the need for a better understanding of age-related variance depicted by each of these modalities. In general, integrating multiple modalities improves the accuracy of age prediction ^26^, particularly when anatomical information is paired with functional information based on a range of phenotypes ^27^. While such complex models involving multiple modalities or models with complex feature input ^29^ may be able to depict more subtle underlying associations between phenotypic and target variables ^13,14^, the interpretability or identification of brain-age interactions is often impeded. Also, more accurate age predictions do not necessarily align with clinical utility ^30,31^, and guidelines on which set of features best to select are missing ^14^. Various neural markers that reflect the amplitude, phase, peaks, or complexity of M/EEG signals have been used to describe the developing or aging brain. Yet, it remains unclear which ones should be favored. Which aspects of neural activity are most indicative of aging processes? And which may be relevant for individual health and risk assessment?

To address these questions, the present study sets out to first evaluate the predictive value of known neural markers used in lifespan research and, second, explore whether age prediction can be more accurate with novel features that have not previously been used in this context. Specifically, we used the currently largest MEG resting-state dataset in adults aged 18 to 88 years from the Cambridge Centre for Aging and Neuroscience (Cam-CAN) and performed extensive extraction of signal characteristics using highly comparative time-series analyses (hctsa) ^32,33^. The hctsa toolbox enables the computation of a comprehensive collection of features relevant across scientific disciplines. Among others, hctsa includes operations for correlational and distributional measures, measures from information theory, time-series model fitting, forecasting, and fluctuation analysis, Fourier and wavelet transform, basic statistics and trend analyses, as well as statistics from biomedical signal processing ^32^. Considering such an extensive feature space has improved, for example, the classification of brain states compared to traditional measures ^34^ or identified features well-reflective of cortical micro-architecture ^35^. Here, we employed spatially informative prediction models using partial least squares regression for each of these hctsa features, as aging affects distributed areas of the brain and yields better performance than anatomically naive models ^27^.

Our findings reveal substantial differences in the predictive value of signal features and suggest that simple linear time-series measures show promise for robustly capturing lifespan patterns, surpassing traditional spectral features. Further investigation into these novel metrics could offer valuable insights into brain development and aging processes.

## 2. Results

Individual MEG time-series were projected to parcellations of the Schaefer atlas ^36^ using beamforming methods (LCMV) ^37^ and the Fieldtrip toolbox ^38^. We characterized five minutes of cleaned resting-state eyes-closed data for each parcel of each individual using features provided by the hctsa toolbox (n = 5961), conventional, and other custom features (n = 26, see methods section for details). The set of conventional features entailed frequency-specific power, amplitude- and phase-based connectivity, the 1/f exponent (slope), and alpha peak frequency (instantaneous alpha peak frequency, center of gravity, see methods section for details). For each feature, the values across all parcels were used to predict the individual age of participants (n = 350, 18-88 years) using partial least squares regression (PLSR) and 10-fold cross-validation (50 repetitions). We evaluated model performances with the correlation between the real and predicted age (Pearson’s r), mean absolute error (MAE), and the predicted R^2^.

### 2.1. Performance of conventional MEG measures

Correlations between chronological and predicted age for conventional features yielded results between r = 0.17 (MAE = 17.03, std = 1.51, R^2^ = -0.13) and r = 0.65 (MAE = 11.93, std = 1.39, R^2^ = 0.42), with the lowest performances of amplitude- and phase-coupling measures and best performances of measures depicting alpha peak frequencies (**Figure 1A**). To evaluate whether the prediction models performed better than chance level, we set up a dummy model by applying the mean age of the training data in each fold of the cross-validation procedure. The cross-validation splits were kept constant across features. Hence, the same dummy model served as our baseline for each feature with an MAE of 16.78 years (std = 0.44, R^2^ = 0, prc_2.5,97.5_ = [15.91 17.65]). The top-ranking conventional measure was center of gravity (r = 0.65, R^2^ = 0.42, **Figure 1B**, **Supplementary Figure 1**), a measure of alpha peak frequency, exceeding the generalization error of the dummy model by about -4.85 years (std = 1.44, prc_2.5,97.5_ = [-7.11 - 2.60]), closely followed by instantaneous alpha peak frequency (MAE diff = -4.36, std = 1.49, prc_2.5,97.5_ = [-6.78 -2.00]). Age predictions using original, aperiodic-adjusted alpha power or the 1/f slope were also more accurate than chance-level prediction by about ∼3.5 years (percentiles ranging from prc_2.5_ = -5.83 to prc_97.5_ = -1.02; **Supplementary Table 1**). Measures characterizing signal power in the theta, gamma, and beta frequency bands showed about 3 to 1.5 years difference to the dummy model (percentiles ranging from prc_2.5_ = -5.07 to prc_97.5_ = 0.45; **Supplementary Table 1**). Connectivity measures generally performed worse than original and adjusted power in most of the frequency bands with 2.3 years or no improvement over the dummy model (percentiles ranging from prc_2.5_ = -4.42 to prc_97.5_ = 2.79; **Supplementary Table 1**). In general, pair-wise differences for the mean absolute error were highly stable across folds and repetitions for most of the conventional measures, with 76 to 100 % better performance of the true than the dummy models. This ratio dropped to 67 – 46 % for phase-connectivity (dwPLI) in the theta, beta, and gamma ranges and for amplitude-coupling in the gamma band. Overall, these findings suggest remarkable differences among conventional MEG signal characteristics in their ability to predict age based on resting-state data.

**Figure 1.**
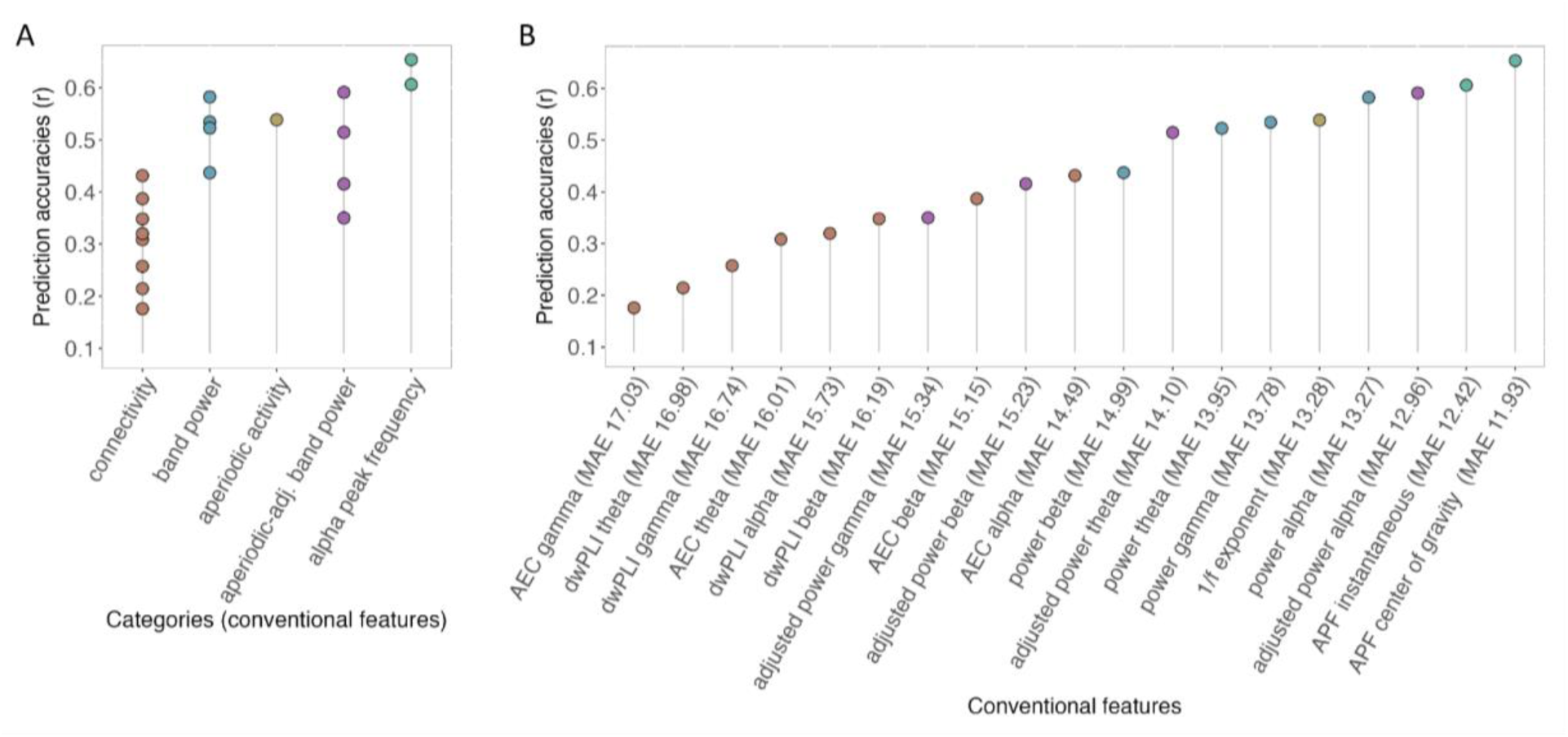
Prediction accuracies of conventional features. Colored dots indicate the predictive value of commonly studied metrics quantified by the correlation (Pearson’s r) between true and predicted age across adulthood (n = 350, 18-88 years). **A)** summarizes the conventional features within their categories and **B)** indicates single feature performance. Adjusted power refers to aperiodic-adjusted power as computed using spectral parameterization methods ^39^. Abbreviations: AEC = amplitude envelope correlation, dwPLI = debiased weighted phase-lag index, APF = alpha peak frequency, MAE = mean absolute error.

### 2.2. Performance of novel time-series measures

Next, we explored whether other MEG phenotypes, previously not considered in developmental and aging neuroscience, would be more sensitive to lifespan variability. We analyzed all time-series features available from the hctsa toolbox that passed the filtering procedure and the custom set (see *Methods*) leaving 5968 features for further probing. In the same cross-validation loop as the conventional measures, we fitted PLS regression models at the level of each feature and all brain parcels. Features that outperformed conventional ones, hence exceeding the prediction accuracy of the top feature peak alpha frequency, will be highlighted in the following (detailed results are listed in **Supplementary Table 1**). 113 novel measures had a higher prediction accuracy than r = 0.7 (MAE = 11.07 to 10.34; std = 1.33 to 1.14; R^2^ = 0.48 to 0.56) and showed an improvement of 5.71 years (std = 1.37, prc_2.5_ = -7.81 to prc_97.5_ = -3.55) to 6.44 years (std = 1.21, prc_2.5_ = -8.23 to prc_97.5_ = -4.39) compared to the chance level prediction based on the mean age of the training data. Figure 2 summarizes the features of interest within their respective categories. Among others, top-performing features included coefficients of autocorrelation, autoregression, wavelet decomposition of time-series, and scaling parameters such as the Hurst exponent or fluctuations. Furthermore, distributional measures sensitive to local extrema or stationarity of the signal or time-series forecasting have been indicative of aging.

**Figure 2.**
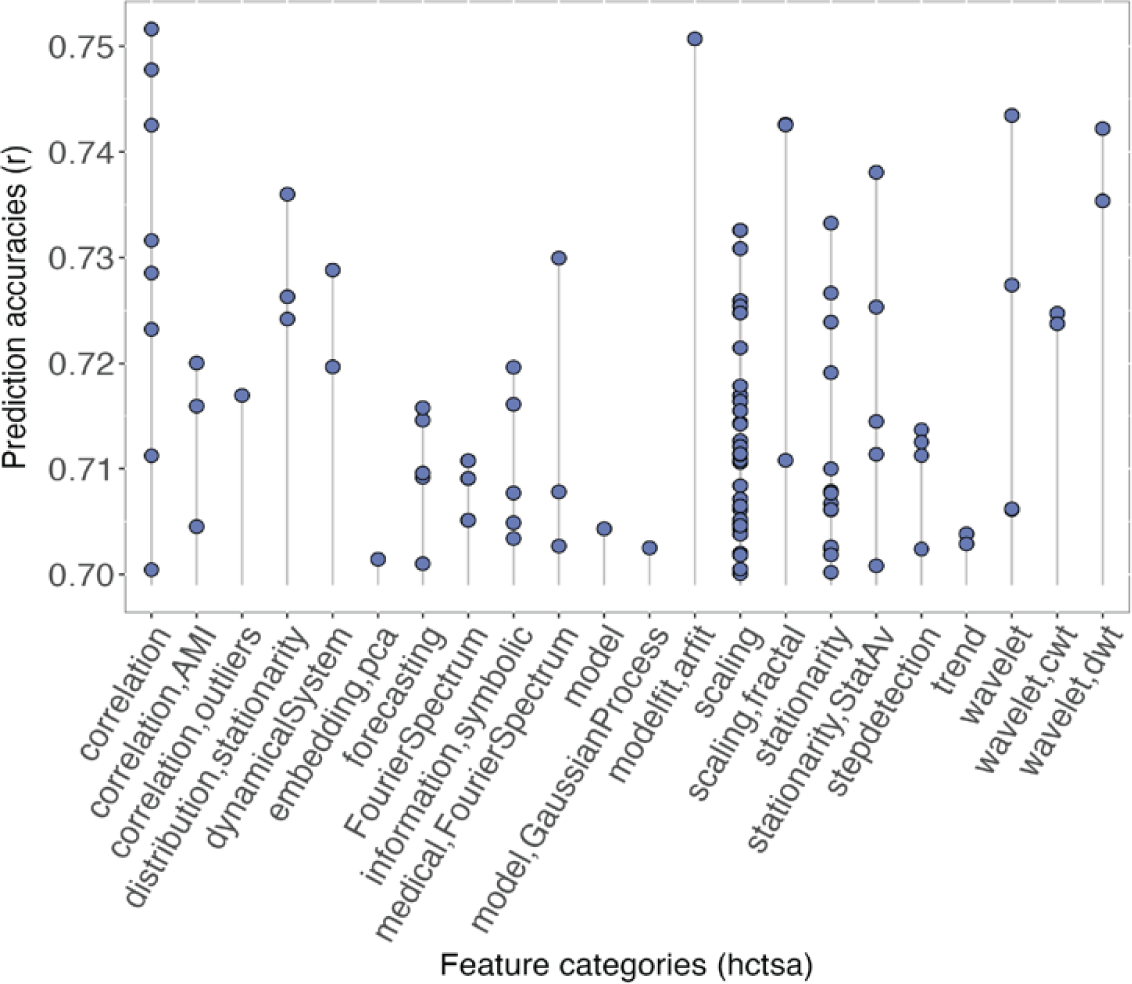
Prediction accuracies of novel features. Blue dots indicate the predictive value (Pearson’s r) of single features computed using highly comparative time-series analysis (hctsa) with higher accuracies (r > 0.7) than conventional features shown in Figure 1. The labels show operational categories to which the investigated features belong.

Of note, many relevant features are correlated, because they cover similar parameters within the same feature category and/or reflect the same aging processes. For example, autocorrelation at time lag 11 was most accurate in predicting age, but other lags (9-15) performed similarly well (**Supplementary Figure 2**), indicating that the function per se captures age-related variability. **Supplementary Figure 3** illustrates how the values of the top features across subjects relate to each other. To elucidate the underlying regional patterns, we transformed the PLS regression weights according to Haufe and colleagues ^40^. This transformation allows the interpretation of the beta weights in terms of a linear forward model. Spatial clustering of the transformed prediction weights for all well-performing features (r > 0.7) revealed patterns that can be grouped according to cortical and subcortical organization (Figure 3). Around four clusters explained most of the variance in the prediction patterns across these features, including the visual cortex and the temporal-parietal cortex with hippocampal structures, putamen, and pallidum. Two other clusters encompassed frontal-parietal areas and frontal lobe structures together with subcortical nuclei and the hippocampus. These findings suggest that these features reflect aging in biologically plausible brain networks.

**Figure 3.**
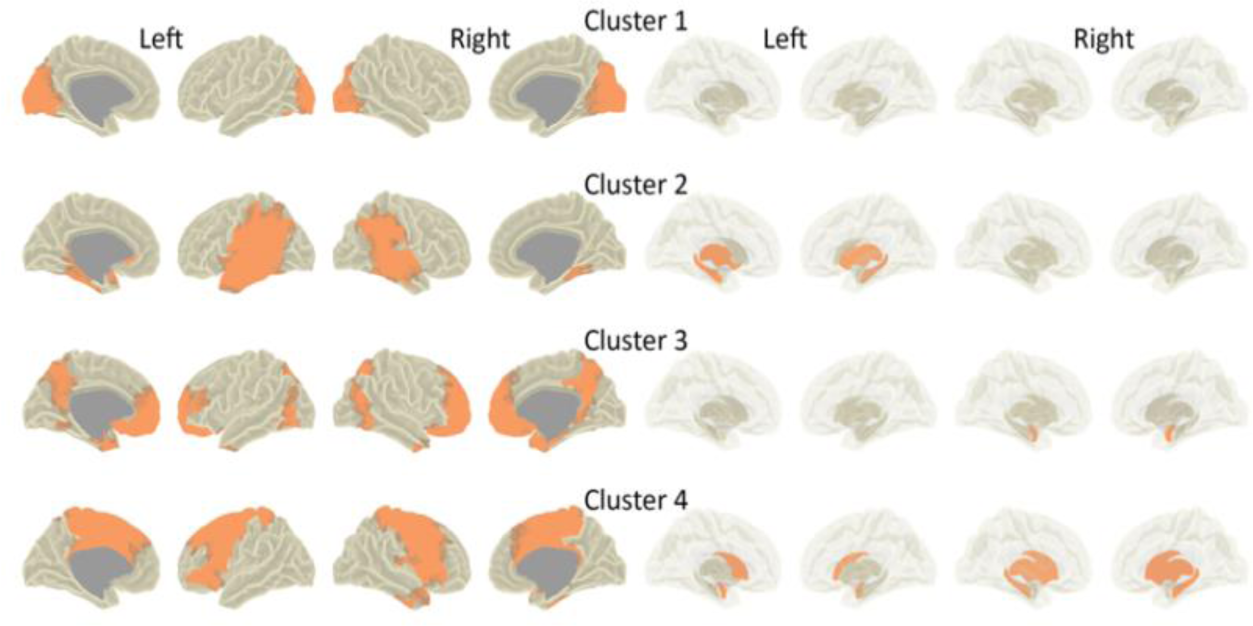
Regional patterns of features predictive of age. Shown are four clusters that represent the prediction weights of highly accurate features (r > 0.7, n = 113) and point to age-related variability in biologically plausible brain networks. PLSR weights for each feature were transformed ^40^ and z-scored before clustering. Left panel = cortical regions, right panel = subcortical regions including the hippocampus. Left = left hemisphere, right = right hemisphere.

### 2.3. Autocorrelation indicates lifespan variability

Autocorrelation at a time-lag of 11 was the feature that predicted age-related variability best among all novel features investigated. Autocorrelation measures the correlation between a time-series and a time-shifted version of itself. Given the sampling frequency of 300 Hz in our study, the autocorrelation at lag 11 corresponds to a time delay of 36 ms and predicted age with an accuracy of r = 0.75 (MAE = 10.34, std = 1.14, R2 = 0.56, Figure 4A). As shown in Figure 4B, positive, transformed prediction weights were present in occipital brain regions and indicated an increase in autocorrelation from young to old age. In other words, self-similarity between MEG signals over time was present at a higher degree in elderly people than in young individuals. For a more comprehensive understanding, we plotted the autocorrelation function up to a time-lag of 133 ms in a brain parcel with positive weights as part of the visual cortex (Figure 4C **lower**). The function peaks around lag 30, suggesting that the MEG signal exhibits a repeating pattern with a period of about ∼100 ms or 10 Hz. A shift in the peak of the autocorrelation function can be noted for the different age groups, hence, a shift in the dominant frequency with age is evident. For comparison, we also plotted the autocorrelation function for a brain parcel of the temporal cortex with negative transformed weights (Figure 4C **upper)**. Here, the function was much flatter in shape, and at a time-delay of 36 ms, a reverse relationship, that is, a decrease of autocorrelation with age was observed, mainly for the young versus the middle and old age groups. In general, the age prediction improved by up to 1.5 years when using autocorrelation of MEG time-series at this lag rather than peak alpha frequency as the best-performing conventional metric. As such, autocorrelation measures seem to contain valuable information about age-related changes in neural activity.

**Figure 4.**
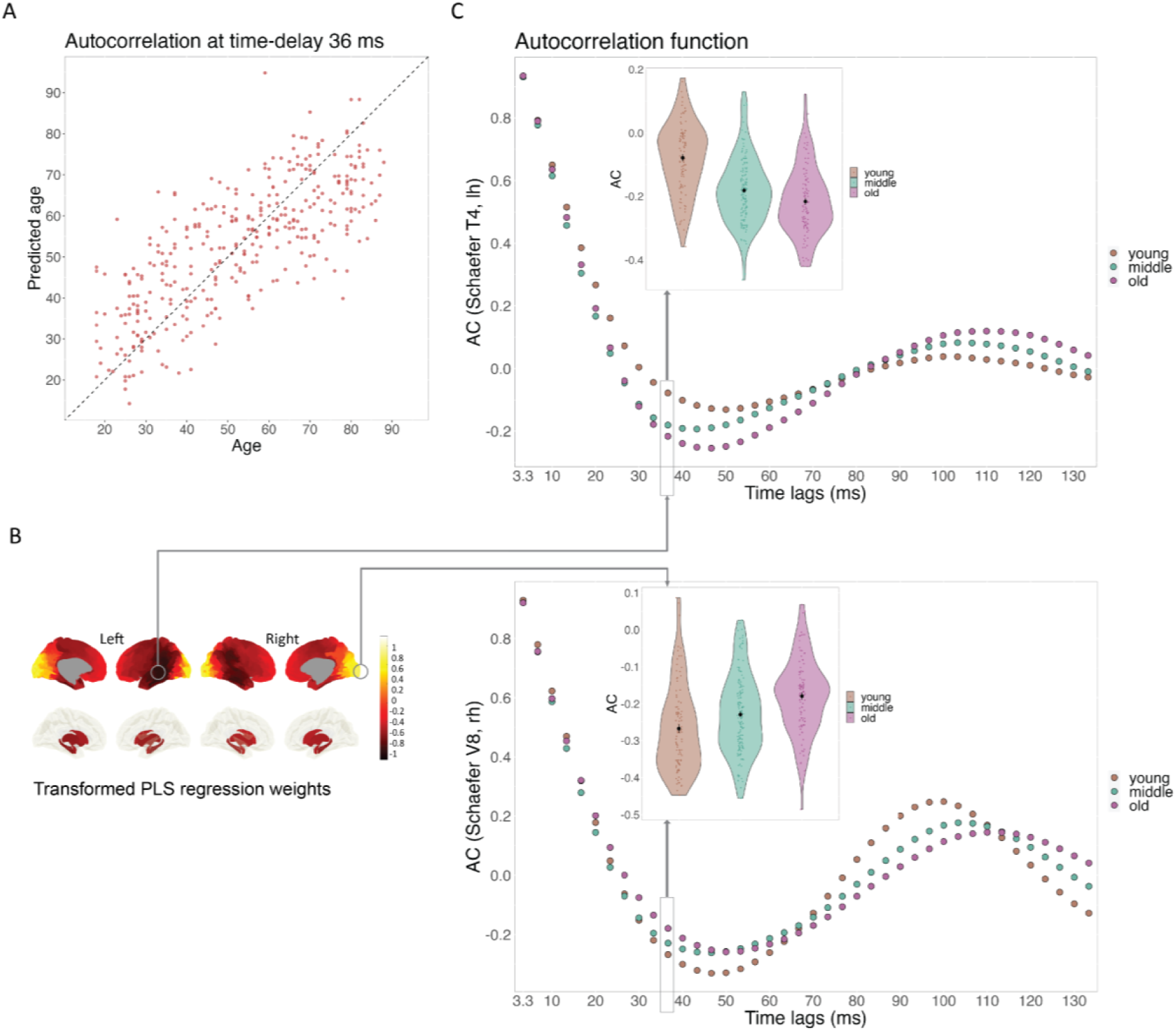
Age prediction using temporal autocorrelation. **A)** The scatterplot illustrates the correlation between true and predicted age when autocorrelation (AC) at a time-delay of 36 ms (lag 11) was used for prediction (r = 0.75, MAE = 10.34). **B)** Plotted are the Haufe-transformed prediction weights of AC lag 11 for cortical (top) and subcortical regions including hippocampal structures (bottom). Yellow colors reflect positive weights and dark colors represent negative weights. Left = left hemisphere, right = right hemisphere. **C)** The upper panel depicts the autocorrelation function and values for a brain parcel (Schaefer T4) in the temporal cortex, indicating decreased autocorrelation in the middle to old age compared to the young age. In the lower panel, the autocorrelation function is plotted for a brain parcel in the visual cortex (Schaefer V8) for each time-lag and averaged for each age group. Here, old individuals had higher values than young individuals at lag 11. Black dots indicate the mean of each age group (*n*_young_ = 100, *n*_middle_ = 150, *n*_old_ = 100) and colored dots represent individual data points. Age groups: young = 18-38 years, middle = 39-68 years, old = 69-88 years.

### 2.4. Confounder analysis

We further examined if other variables than age confounded the prediction analyses, as out-of-interest variables can hamper the validity and generalizability of prediction models ^41^. Specifically, we assessed if sex or estimated total intracranial volume (eTIV) might have driven the age predictions using the time-series features. After corrections for multiple comparisons, there was no confounding bias of sex or eTIV for any of the investigated features (p _FDR_ > 0.05). Hence, these two variables were not more strongly related to the predicted age values than what can be expected for the association with observed age. Detailed results are reported in **Supplementary Table 2**.

## 3. Discussion

We provide a comprehensive overview of the predictive value of neural metrics for age across adulthood by employing prediction and cross-validation techniques to minimize the overfitting problem. The extensive profiling of regional MEG signals during rest in a large adult sample unveiled notable differences between metrics traditionally used in the context of development and aging neuroscience. Intriguingly, other, less considered time-series features reflected age-related changes in neural dynamics at the milliseconds scale more robustly. We propose that simple linear measures such as autocorrelation show substantial promise to offer valuable insights into brain development and aging.

So far, lifespan studies directly comparing the performance of single MEG/EEG markers have been rare. Previous machine learning studies point to a relevant role of regional alpha and beta frequency power for age prediction ^27^, which is in line with other prediction work using unimodal EEG ^42^. We confirm these findings by demonstrating that alpha power carries more predictive information than power in other frequency bands or even connectivity measures for the adult lifespan. Crucially, our study adds that the shift of the alpha peak with age tends to be a more robust measure than spectral power. A shift in the alpha peak is a known developmental phenomenon ^43–45^ such that children have a peak frequency at around 6 Hz, which increases up to adolescence, potentially masking age effects in alpha power when using fixed frequency limits ^46^. In contrast, a global slowing of the alpha peak frequency has been reported across the adult lifespan ^47–49^. A global decrease was also present in our sample (compare **Supplementary Figure 1**), but a combination of a decrease in visual-temporal regions and an increase in frontal and motor areas were particularly predictive of age. Interestingly, alpha peak shifts link to cognitive performance such as attention during development ^46^ and executive functions in old age ^50^, and predicted cognitive decline over 10 years when combined with the aperiodic exponent ^51^. Also in our study, the aperiodic exponent ranked comparably high in predictive power, underlining the importance of taking non-rhythmic aspects of neural dynamics into account when studying age.

Inherent to previous age-related studies is that, so far, the focus has mostly been on MEG or EEG spectral signal characteristics that are historically in place and distinguished healthy from pathological brain states. We extend this perspective and suggest focusing additionally on time-related aspects of neural dynamics, such as the self-similarity of the signal as depicted by the autocorrelation function. For autocorrelation, generalization errors ranged from 11 to 10.3 years (R^2^ = 0.5-0.56) for the most predictive time-lags. For comparison, others using MEG CamCAN data have found deviations of approximately 9 years (R^2^ = 0.55, ^26^), ∼7 years ^27^, or ∼ 8 years (R^2^ ∼ 0.7, ^29^) for models that were trained on different combinations of several MEG features. Engemann and colleagues (2022) also considered a set of less conventional signal characteristics besides spectral features, such as Hurst or Hjorth parameters, wavelet decomposition, or fractal dimensions. However, this study focused on accurate age prediction rather than identifying interpretable MEG phenotypes associated with age. Other features from the hctsa toolbox also showed comparably high prediction accuracies as autocorrelation but tended to be more complex and difficult to interpret. The strong correlations among predictive features across individual brains (compare **Supplementary Figure 3**) and the similarity of the prediction patterns, as assessed using clustering, indicate that these features are sensitive to similar neuronal dynamics. Therefore, we further focus on temporal autocorrelation as a robust measure of the adult lifespan.

The autocorrelation function reflects the similarity of the neural signal across time-lagged copies of itself. In our study, high prediction accuracies were achieved based on feature values in the occipital and temporal cortex. Of note, the cycle of the autocorrelation function in the visual cortex peaks at around 10 Hz and therefore corresponds to the well-studied alpha activity. This is plausible given that alpha oscillations are generally dominant in posterior brain areas during an eyes-closed resting-state scenario like in our study. However, signal self-similarity near the first minimum of the function was highly predictive of age, including the time-lags between 30 ms to 50 ms. This is of great interest for two reasons. First, the initial decay of the autocorrelation function is related to aperiodic components in the signal. Second, alpha activity is not sinusoidal, and the autocorrelation at lags other than 100 ms is sensitive to the specific waveform shape of the regional brain activity. Autocorrelation therefore provides a versatile set of features that can capture various aspects of age-related changes in electrophysiological signals. In a similar vein, temporal autocorrelation derived from much slower fMRI signals has been an accurate index of the connectome by explaining changes in complex graph network topologies with age and thus likely reflects the fundamental reconfiguration of the brain’s network during aging ^52^. More generally, the decay constant of the autocorrelation function, also known as neural timescale, is thought to enable the maintenance of information in a hierarchical manner ^53^. That is, timescales are longer in association areas for higher-order tasks, but are shorter in sensory regions coupled to stimuli in the environment ^54–56^. A compression of neuronal timescales with age was observed in most regions except the visual cortex, such that older adults showed faster time scales in intracranial recordings than young adults ^56^, similar to what can be observed in our data.

Notably, the spatial prediction patterns of autocorrelation varied between different time lags (**Suppl Figure 2**), suggesting that a combination of linear changes in autocorrelation at specific spatial configurations may distinguish individuals at different ages. Nevertheless, a reversed pattern of autocorrelation between sensory-motor and association areas emerges, in line with the notion that the brain develops along this axis of brain organization ^57^, which seems observable up into old age. Consistently, brain state dynamics differed between visual areas and higher-order networks ^58^ across the adult lifespan and were linked to cognitive performance, with a reduction in older people being more strongly associated with shifts between these states than in young people. As previously mentioned, age-related changes in the visual cortex have also been reflected in phase-based connectivity across different frequency bands ^4^, but the predictive power of phase-related measures was rather poor in our study and others ^26^. This may be because estimations of the phase of M/EEG signals are generally susceptible to noise, making it a less robust metric than autocorrelation, which is well-normalized between -1 and 1. It is also important to note that we only looked at the averaged connections between brain regions rather than taking within- and between network connections into account thereby reducing potentially valuable, age-specific information. Furthermore, we observed a decrease in autocorrelation in the temporal cortex, particularly in the age groups after 40 years of age. Again, previous studies have identified the temporal cortex as an area associated with strong M/EEG power increases across the adult lifespan ^3,4^, with an increase over the adult lifespan.

However, the signal amplitude is not a normalized measure and strongly varies within and across individuals, hampering the distinction between the age groups and the generalizability of the model across the lifespan. Overall, these functional changes divergent between sensorimotor and association regions are likely related to the microstructural and transcriptomic backbone of the brain ^57^. Intriguingly, neural timescales have also been correlated with biological factors that shape neuronal dynamics, such as genes related to voltage-gated ion transporters and GABA and chloride channels ^56^. These findings highlight new opportunities to unravel the complexity of development and aging mechanisms at the intersection of brain pathology through temporal autocorrelation.

It is important to keep in mind that partial least squares regression, as used here, is a method that detects linear relationships in the data. Hence, algorithms sensitive to non-linear relationships might yield different outcomes for some of the features investigated in this study. Indeed, previous work has observed quadratic age trajectories for MEG characteristics in the beta-gamma range ^3,4,7^. However, machine-learning techniques detecting non-linear variability often yield less interpretable results. Another caveat in our study concerns the present results for deep brain structures. Highly predictive feature weights loaded on subcortical regions and the hippocampus, but these findings should be interpreted with caution. It is debatable to what extent activity from deep brain areas can be reliably detected using MEG or if the observed activities are noise projections through beamforming. Yet, simultaneous MEG and intracranial recordings have demonstrated that signals from the mesial temporal lobe and the thalamus can be read out at the brain surface ^59^. Also, subcortical features derived from MRI have been more reliable age predictors than cortical features ^26^, pointing to informative changes with age presumably affecting the brain at the functional level.

In sum, we demonstrate that the spatial distribution of single, simple, and easily interpretable MEG time series features across the brain is highly predictive of individual age. Further improvements are expected by models that combine features, such as incorporating all autocorrelation lags or different synergistic features. Autocorrelation of non-invasively measured electrophysiological brain activity might serve as a dynamic marker for healthy aging and early intervention, and leverage brain age predictions based on structural properties of the brain. Given the simplicity and interpretability of autocorrelation as a feature, future studies might identify, with the help of biologically inspired generative computational models, the exact anatomical, chemical, or functional properties that change with age.

## 4. Materials and Methods

### 4.1. Data and participants

We used cross-sectional open-access data from the Cambridge Centre for Aging and Neuroscience (Cam-CAN) ^60^. From 650 resting-state MEG data sets and T1-weighted MR images available from phase two of the study ^61^, we further analyzed a subset of cleaned data (*n* = 350) as reported elsewhere in detail ^4^. Our sample included 50 healthy individuals of each age decade between 18 to 88 years and was balanced by sex. All individuals were cognitively normal, without communication problems, major physical impairments, neurological or psychiatric conditions. The study was conducted in accordance with the Declaration of Helsinki ^62^ and approved by the local ethics committee, Cambridgeshire 2 Research Ethics Committee. Data collection and sharing for this project was provided by the Cam-CAN (https://camcan-archive.mrc-cbu.cam.ac.uk/dataaccess/).

### 4.2. MRI acquisition

Anatomical images were acquired using a 3T Siemens TIM Trio scanner with a 32-channel head coil. T1-weighted images were derived from 3D MPRAGE sequences with TR = 2250ms, TE = 2.99ms, TI = 900ms; FA = 9 deg; FOV = 256×240×192 mm; 1 mm isotropic; GRAPPA = 2; TA = 4 mins 32 s), and T2-weighted images from 3D SPACE sequences with TR = 2800 ms, TE = 408 ms, TI = 900 ms; FOV = 256×256×192 mm; 1 mm isotropic; GRAPPA = 2; TA = 4 mins 30 s).

### 4.3. MEG acquisition

Data were recorded using a 306-channel VectorView MEG system (Elekta Neuromag, Helsinki) with 102 magnetometers and 204 planar gradiometers (sampling at 1 kHz with a 0.03 Hz online high pass filter). Individuals were assessed during the resting-state with their eyes closed for 8 min and 40 s in a seated position and a magnetically shielded room at a single site (MRC Cognition and Brain Science Unit, University of Cambridge, UK). The head position within the MEG helmet was continuously measured by four coils. Simultaneously, cardiac signals and eye movements were recorded using electrocardiogram (ECG) and electrooculogram (EOG, horizontal and vertical), respectively.

### 4.4. Individual head models and source projection

T1- and T2-weighted images were reconstructed using FreeSurfer 6.0.0 (https://surfer.nmr.mgh.harvard.edu/) and MEG sensor data further projected to individual cortical surfaces derived from SUMA ^63^. Based on the ‘fsaverage’ template mesh provided by FreeSurfer and SUMA (afni.nimh.nih.gov/download/), cortical surfaces were resampled to 1,002 vertices per hemisphere (ld factor = 10). In addition, we included deep brain structures such as bilateral amygdala, hippocampus, thalamus, caudate, putamen, and pallidum reconstructed from the fsaverage template. Each region was converted to surfaces and spatially normalized to Montreal Neurological Institute space (DARTEL; SPM12; fil. ion.ucl.ac.uk/spm/software/spm12/) using CAT12 DARTEL template (neuro.uni-jena.de/cat/), yielding 2338 brain vertices for each participant as described elsewhere ^64^. The individual meshes were realigned to the Neuromag sensor space based on anatomical landmarks previously defined by Cam-CAN. Leadfields and individual head models were computed using the “single shell” method implemented in Fieldtrip.

### 4.5. MEG processing

Cam-CAN provided preprocessed MEG data through their repository including temporal signal space separation (Elekta Neuromag Maxfilter 2.2, 10s window, 0.98 correlation limit) for external artifact and line noise removal (50 Hz and its harmonics) as well as correction for bad channels and head movements. We then resampled the data to 300 Hz using Fieldtrip ^38^ and, after filtering (first order Butterworth high-pass filter at 1 Hz), segmented the data into segments of 10 s length. Further processing steps are described in detail here ^4^. In short, we followed an automatic approach to identify and reject segments containing artifacts from muscle movements (see https://www.fieldtriptoolbox.org/tutorial/automatic_artifact_rejection/), low-pass filtered the data at 70 Hz (first order Butterworth) and used independent component analysis (ICA) to identify ocular and cardiac artifacts. Components were selected based on their similarity to EOG signals (average coherence > 0.3 and amplitude correlation coefficient > 0.4) and coherence with the recorded ECG signal (average coherence > 0.3) or based on the averaged maximum peaks time-locked to the ECG (QRS complex). The selected components were manually checked for each data set. In a few cases, the relevant components had to be visually identified because the ECG/EOG was noisy. Subsequently, cleaned data was inspected for quality control and vigilance of individuals according to the criteria of the American Academy of Sleep Medicine (https://aasm.org/). Finally, 30 segments with clean and “awake” data were randomly selected as previous work showed good reliability for MEG spectral metrics computed on five minutes of data ^65^. Signals from only the magnetometers (102 channels) were then projected to the source level using unit-noise gain linear constrained minimum variance (LCMV) beamforming ^37^. Spatial filters were constructed (regularization lambda = 5%, free-orientation forward solution) and multiplied with the sensor time series data, resulting in source-level time series at 200 parcels of the Schaefer atlas ^36^.

### 4.6. Characterization of projected time-series

We derived 7525 features from the hctsa toolbox ^32^ to characterize the projected signals at each brain parcel across age. After removing features that produced constant or complex values, outliers, or invalid numbers, a total of 5961 hctsa features were kept for further analysis. Moreover, we added other features that we deemed important due to their application in the signal processing literature such as the Hjorth parameters (activity, mobility, complexity ^66^), mean absolute deviation, zero crossing, zero crossing derivative, and weighted permutation entropy ^67^. To compare the predictive performance of this explorative set of features, we computed conventional features that have been extensively applied in the literature of cognitive and clinical neurosciences including frequency-specific amplitude-^68^ and phase-coupling (debiased weighted phase-lag index ^69^), absolute power, and measures of alpha peak frequency (center of gravity ^70^, local maxima, instantaneous alpha peak frequency ^71^). Due to estimation problems of the alpha peak frequency using local maxima in fronto-central regions, this metric was dropped. Power and connectivity measures were estimated in four frequency bands, i.e., theta (4-8 Hz), alpha (8-13 Hz), beta (13-30 Hz) and gamma (30-60 Hz). We also passed the source power spectrum (1-60 Hz) to the specparam algorithm ^39^ with the following settings: peak width limits [1-8]; maximum number of peaks: infinitive; minimum peak height: 0; and aperiodic mode: fixed. This resulted in an aperiodic component (1/f exponent or slope) and periodic (“oscillatory”) components, here referred to as adjusted power.

### 4.7. Prediction analysis

Individual age values were predicted from whole-brain parcellations using partial least squares regression for *n* = 5987 features and stratified 10-fold cross-validation. In each fold or iteration, the entire sample (*n* = 350) was split into nine train sets and one test set containing approximately the same age distribution. The test set was subsequently used to predict age from the unseen data. This procedure was repeated 50 times, and the resulting parameters were averaged to increase the robustness of the results. Partial least squares regression (PLSR) is a multivariate, linear technique that predicts one set of data from another ^72^, in our case individual age from MEG feature values at cortical and subcortical parcels. Based on singular value decomposition, PLSR iteratively finds orthogonal latent components of the predictors that correlate best with the response variable (age), producing corresponding regression weights for each predictor, i.e., brain parcel. We transformed these weights according to Haufe and colleagues ^73^ for the interpretability of the resulting topographical maps. As the estimated latent components of the prediction model are uncorrelated, this can be obtained by the covariance of the observed data (participants x parcels) multiplied by the PLSR weights. This transformation makes the PLSR output interpretable in terms of a forward model ^40^. All analyses were performed using Matlab (R2018b) and for each of the conventional and novel features, respectively. We assessed the predictive power of the features using the correlation between the real and predicted age (Pearson’s r), the mean absolute error (MAE) in years, and the prediction R^2^ ^74,75^. Furthermore, we created a dummy model to test whether the models of interest performed better than the chance level, and against which generalization errors of all models can be compared. We used the mean age of the training data in each cross-validation fold, resulting in an MAE of 16.78 years (std = 0.44, R^2^ = 0, prc_2.5,97.5_ = [15.91 17.65]).

### 4.8. Confounder analysis

We assessed if sex and total intracranial volume have potentially confounded the prediction analyses as these variables have contributed to variance in spectral MEG features in previous work (e.g. ^3,4,76,77^). We implemented a model-agnostic approach using the partial confounder test from the mlconfound package (https://mlconfound.readthedocs.io) that tests the conditional (in-)dependence between the predicted values and each potential confounder ^41,78^. If significant, the predicted age values are more strongly associated with the confounding variable than what is expected from the association between the observed age and the confounders. P-values were computed based on the defaults (1000 permutations, Markov-chain Monte Carlo process set to 50) and FDR-corrected across all features tested (*n* = 5987).

### Data and code availability

Raw data were provided by the Cam-CAN project and are available at https://camcan-archive.mrc-cbu.cam.ac.uk/dataaccess/underspecifiedconditions. Single-subject feature values can be downloaded from https://osf.io/h43mz/. All code used for the preprocessing, statistical analyses, and visualization of the results have been deposited in https://github.com/chstier/Aging_timefeats.

## Acknowledgments

Data collection and sharing for this project was provided by the Cambridge Center for Ageing and Neuroscience (CamCAN). Cam-CAN funding was provided by the UK Biotechnology and Biological Sciences Research Council (grant number BB/H008217/1), together with support from the UK Medical Research Council and the University of Cambridge, UK. This work was further supported by the German Research Foundation (DFG; GR 2024/11-1 to JG; FO 750/5– 1 to N.K.F.). We acknowledge support by the Open Access Publication Funds of the University of Münster. We are grateful to Professor Ben Fulcher for valuable discussions on the methods.

## Author contributions

Conceptualization J.G., C.S.

Methodology J.G., C.S., E.B., J.F., N.F., A.W.

Investigation C.S., J.G.

Writing – original draft C.S.

Writing – review & editing E.B., J.F., N.F., A.W., U.D., J.G.

Visualization C.S.

Funding acquisition N.F., J.G.

## Supplementary Figures

**Suppl Figure 1.**
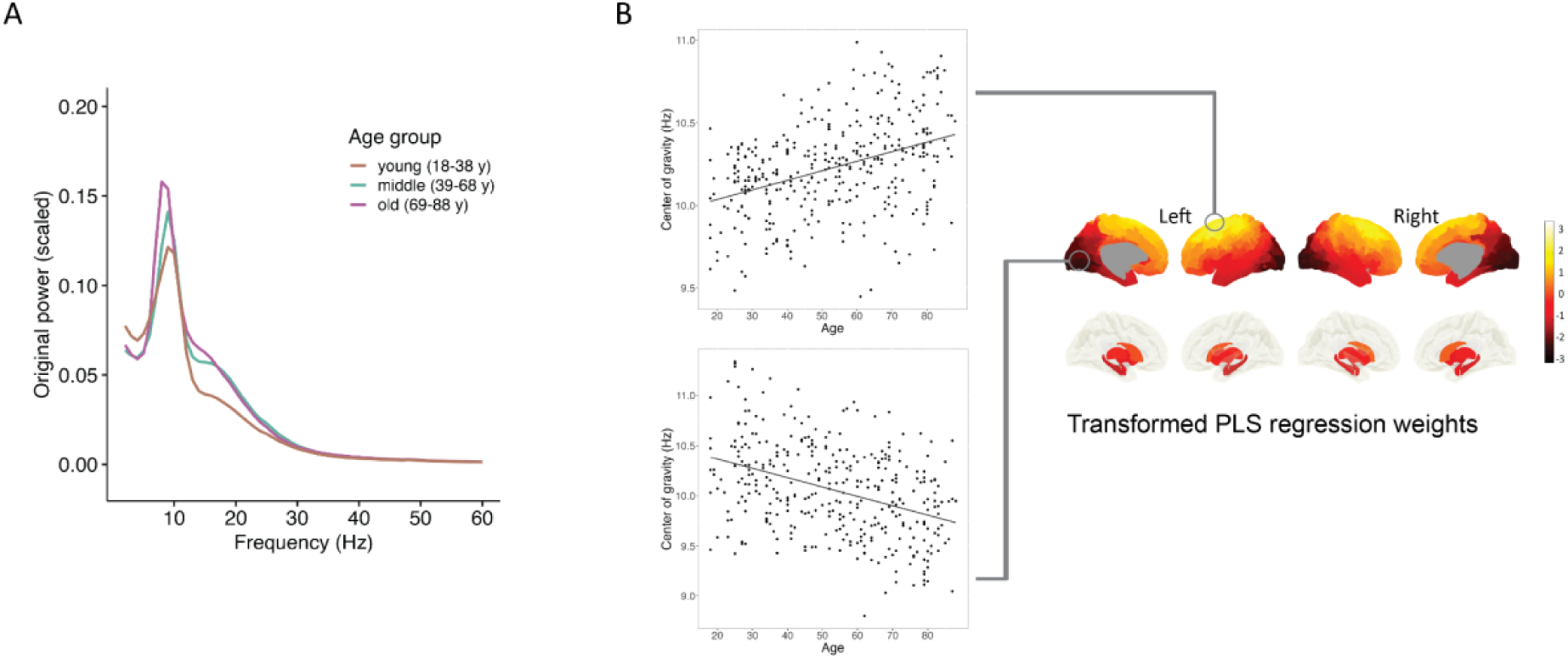
Power spectrum and age prediction results for the center of gravity (alpha band) **A)** Power spectrum (absolute power in femtotesla, scaled) averaged across all brain parcels and for three age groups: young = 18-38 years (n = 100), middle = 39-68 years (n = 150), old = 69-88 years (n = 100) **B)** Plotted are the Haufe-transformed prediction weights of center of gravity for cortical (top) and subcortical regions including hippocampal structures (bottom). Yellow colors reflect positive weights and dark colors represent negative weights. The scatterplots show individual data points and linear associations with center of gravity in the respective brain parcels. Left = left hemisphere, right = right hemisphere.

**Suppl Figure 2.**
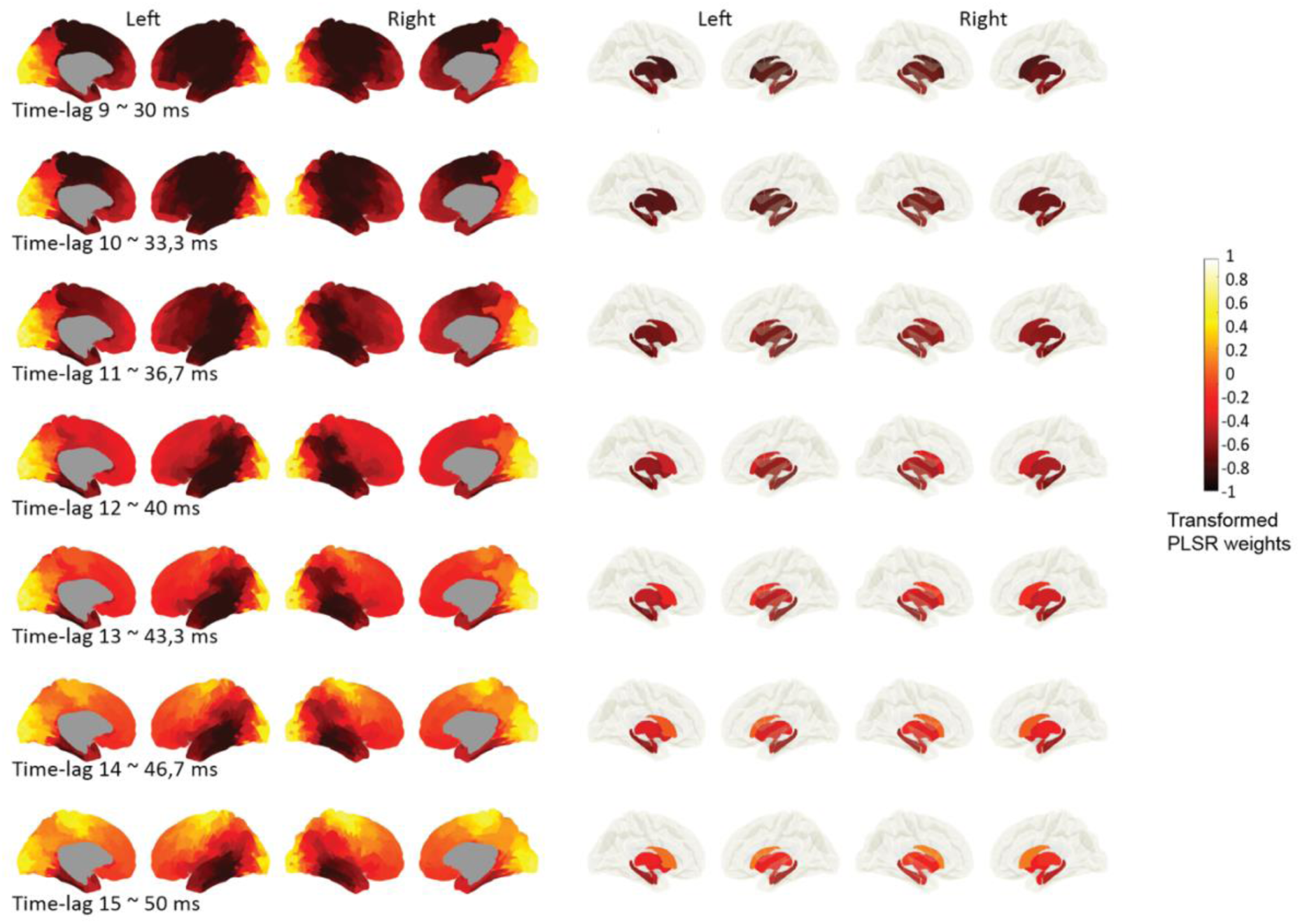
Temporal autocorrelation at highly predictive time-lags. This plot indicates Haufe-transformed prediction weights of autocorrelation at different time lags. Prediction performances were all above r > 7, i.e., lag 9: r = 0.73 (MAE = 10.73); lag 10: r = 0.75 (MAE = 10.45), lag 11: r = 0.75 (MAE = 10.34), lag 12: r = 0.74 (MAE = 10.57), lag 13: r = 0.73 (MAE = 10.75), lag 14: r = 0.72 (MAE = 10.82), lag 15: r = 0.71 (MAE = 11.04). Shown are cortical (left panel) and subcortical regions including hippocampal structures (right panel).

**Suppl Figure 3.**
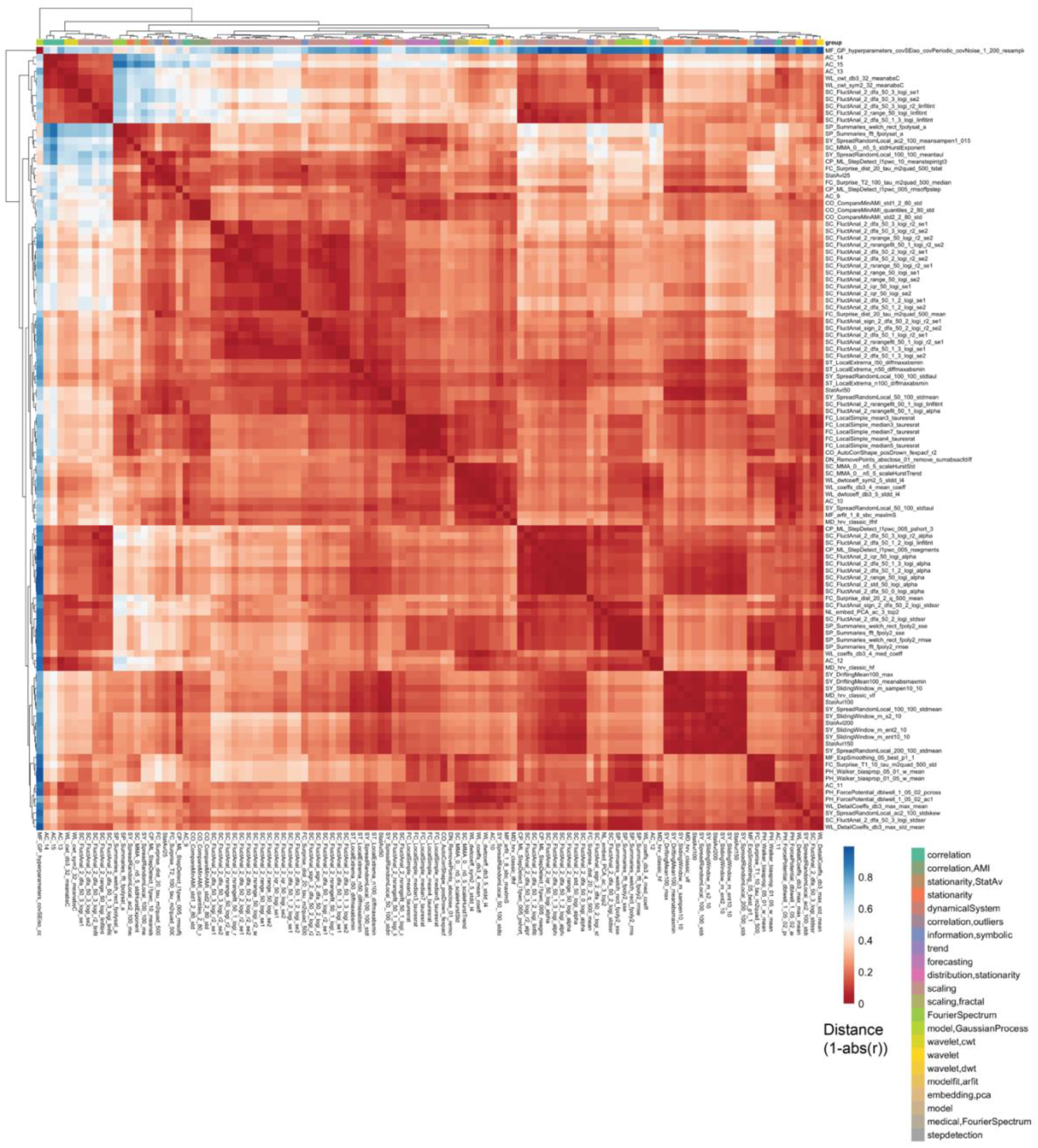
Distance matrix for highly predictive hctsa features (r > 0.7) The matrix highlights the dependency of hctsa features predictive of age. Individual values for each feature (n = 113) across brain parcels were concatenated and correlated. Red colors represent high correspondence among features (1-absolute value of r) and blue colors represent low correspondence. Each row or column indicates a single feature, summarized in their respective feature category (right legend, mixed colors).

